# Morphology of immatures of the thelytokous ant, *Monomorium triviale* Wheeler (Formicidae: Myrmicinae: Solenopsidini) with descriptions of the extraordinary last-instar queen larvae

**DOI:** 10.1101/2021.11.07.467650

**Authors:** Naoto Idogawa, Ayako Gotoh, Shigeto Dobata

## Abstract

The ant genus *Monomorium* is one of the most species-rich but taxonomically problematic groups in the hyperdiverse subfamily Myrmicinae. An East Asian species, *M. triviale* Wheeler, produces both reproductive queens and sterile workers via obligate thelytokous parthenogenesis. Here, we describe the immature forms of *M. triviale* based on light and scanning electron microscopy observations, with a note on the striking caste dimorphism in the last larval instar. The last-instar queen larvae were easily recognized by their large size, “aphaenogastroid” body shape, and rows of doorknob-like tubercles on the lateral and dorsal body surface. This type of queen-specific structure has not been found in ants in general, let alone congeneric species found in Japan. In stark contrast to the queen larvae, worker larvae showed a “pheidoloid” body shape and a body surface similar to other ants. The worker larvae were estimated to have three instars, consistent with previously described congeners. The pupae of both castes had no cocoon, a characteristic commonly described in other Myrmicinae species. In total, the developmental period from egg to adult worker averaged 59 days under 25°C. We discuss possible functions of the tubercles of queen larvae based on previous studies.

## Introduction

Ants (Hymenoptera: Formicidae) are characterized by the division of labor between two phenotypes of females: reproductive queens and sterile workers (Hölldobler & Wilson, 1990). These two castes share the same genetic background and diverge epigenetically in response to socio-environmental factors during immature stages (Rajakumar *et al*. 2018). Furthermore, ant larvae play an important role in colony dynamics and physiological integration by digesting food and distributing nutrients to their nestmates (Dussutour & Simpson 2009; Schultner *et al*. 2017; Masuko 2019). Such behavior emphasizes the importance of morphological study on the immature stages of ants. However, our knowledge of the immature stages of ants, including number of larval instars, growth rate, and developmental periods is surprisingly limited. Even for the worker caste, complete larval descriptions are available in only 0.5% of >13,000 extant species (Bharti *et al*. 2019; Bolton 2021). The reproductive castes, i.e., queens and males, are usually produced in small numbers during limited seasons, meaning there is even less information on their immature stages.

Here, we provide a detailed description of the immature stages of a thelytokous ant, *Monomorium triviale* Wheeler, with a note on their preimaginal caste dimorphism. *Monomorium* Mayr is a globally distributed and speciose genus in the hyperdiverse subfamily Myrmicinae. Although many species have been transferred to other genera by recent phylogenetic studies (Ward *et al*. 2014; Sparks *et al*. 2019), approximately 300 species have been described and a large number of species have yet to be described (Pontieri & Linksvayer 2019). In this genus, the larval morphology has been described from 11 species (Wheeler & Wheeler 1955, 1973, 1980; Bernard 1955; Ettershank 1965), but there is no information on the number of larval instars except for well-studied tramp species, the pharaoh ant, *M. pharaonis* L. (Alvares *et al*. 1993; Pontieri *et al*. bioRxiv) and the flower ant, *M. floricola* Jerdon (Solis *et al*. 2010a). The East Asian species, *M*. *triviale,* is one of the recently found parthenogenetic *Monomorium* species (Gotoh *et al*. 2012; Idogawa *et al*. 2021; Ito *et al*. 2021). Males of *M. triviale* have never been reported, and virgin queens produce both workers and next-generation queens via thelytoky.

In the present study, the number of larval instars of *M. triviale* was estimated based on the distributional pattern of body size. The morphological features of each larval instar were also investigated with light and scanning electron microscopy. Additionally, histological observation was performed on the newly discovered queen-specific structure. Finally, the developmental period required for each stage was examined by rearing experiments.

## Materials and Methods

### Collection of Samples

The nests of *M. triviale* were collected in a thicket consisting mostly of deciduous broad-leaved trees located in the northern suburbs of Kyoto, Japan (35°03’33” N, 135°47’01” E, alt. 90 m) from 2017 to 2021. To obtain all stages of immatures, the nests were transferred into artificial nests in the laboratory and were kept at 25°C.

### Determination of Larval Instars

A number of previous studies on ant larvae have estimated the number of larval instars from the size distribution pattern of head capsule width (e.g., Parra & Haddad 1989). However, Masuko (2017) points out that size-based estimations can overlook the instars with low abundance (especially the first instar). In the genus *Monomorium*, the shape and distribution of hairs have been reported to be useful for the identification of larval instars (Solis *et al*. 2010a). Therefore, we estimated the number of larval instars in *M. triviale* using a combination of morphometrical and chaetotaxical characteristics.

To bracket the lower and upper growth limit of *M. triviale* larvae, we explicitly identified the individuals belonging to the first and last instar. Sixty-two larvae with egg chorion (i.e., just after hatching = definitely first instar) were collected from the egg piles in the nests. Thirty-five prepupae (i.e., just before pupation = definitely last instar) were also collected. We photographed these larvae under a digital microscope (VHX-900; Keyence) and measured the maximum head capsule width using ImageJ software (Schneider *et al*. 2012). Then, we measured 582 randomly chosen larvae and plotted the distributional pattern of the head width in a histogram. The distinct peaks of the histogram were detected to find the intermediate instars between the first and last instar. At least five individuals representative of each peak were extracted and morphological differences such as body hair number and type were examined (see next section). Finally, the number of larval instars was determined on the consensus of qualitative and quantitative traits. The data calculation and visualization was performed with R ver. 4.0.0 software (R Core Team 2021).

During this experiment, we found larvae that were clearly larger than typical last instar worker larvae. These larvae were reared in the laboratory until eclosion and confirmed to be the last instar queen larvae (approx. 50 individuals). We were unable to estimate the number of larval instars for queens in the above manner because *M. triviale* colonies produce only a very small number of queen larvae (1–10 individuals) during the early summer.

### Observation of the immature stages

To characterize the immature stages, the external morphology was investigated with a binocular microscope (SZ40; OLYMPUS Optical, Tokyo, Japan), digital microscope (VHX-900; Keyence, Osaka, Japan), and scanning electron microscope (SEM: VE-8800; Keyence, Osaka, Japan). For the binocular and digital microscope observations, living immatures were carefully fixed on the glass slides using sticky anti-slip gel pads (Seiwa-pro, Osaka, Japan). For the SEM observation, fresh samples were frozen in a deep freezer at −20°C for 15 min then quickly mounted on the aluminum stage with conductive carbon double-sided tape. All observations and photography were performed as soon as possible after sample preparation. Voucher specimens of eggs, larvae, pupae, and nestmate imagos were deposited in the Laboratory of Insect Ecology, Kyoto University, Kyoto, Japan. All terminology used in our larval descriptions follows Wheeler and Wheeler (1976) and Solis *et al*. (2010a) and measures are given as mean ± SD followed by the number (*n*) of observations. For comparison of caste dimorphism, three *Monomorium* species available in Japan, *M. intrudens* Smith, *M. chinense* Santschi, and *M. hiten* Terayama, were additionally observed in the same manner.

Since we found caste dimorphism in the external morphology of the last instar larvae, we examined the histological features of the queen larvae. The larvae were fixed in FAA (pure ethanol:formaldehyde:acetic acid = 16:6:1) for 24 hours and were dehydrated in a graded ethanol series before embedding in paraffin. They were cut at 4 μm and stained with hematoxylin and eosin as described in Gotoh *et al*. (2016). Histological observations were performed with Leica DM IL LED inverted contrasting microscope and Leica MC 170 HD camera (Leica Microsystems, Wetzlar, Germany).

### Developmental Periods

Rearing experiments were performed to determine the developmental periods of each immature stage of *M. triviale*. Due to the difficulty in distinguishing between first and second instar larvae, these stages are grouped as “young larvae”. From the 18 field-collected source nests, a single queen and 10 nestmate workers were isolated in an artificial nest for determination of the egg period. The newly oviposited eggs and hatched larvae were counted daily for approximately 50 days.

For determination of the larval and pupal periods, 10 immatures just before the target stage (i.e., eggs for young larvae period, young larvae for last instar larvae period, last instar larvae for pupae period) were isolated with 10 workers. The numbers of individuals belonging to each stage were counted every day. From the 11 field-collected source nests, 7 artificial nests were prepared for young larvae, 8 nests for last instar larvae, and 7 nests for pupae. Observations were continued for a maximum of one month until all individuals molted to the next stage or died. All experimental nests were isolated in the plastic container (36 × 36 × 14 mm) with gypsum on the bottom and were kept in the laboratory at 25°C. The water and food (mealworms, *Tenebrio molitor* L. cut into approximately 5 mm length) was replenished every 3 days.

## Results

### Determination of Larval Instars

The head capsule width of larvae just after hatching (i.e., definitely first instar; mean ± SD = 0.123 ± 0.007 mm, *n* = 62) and prepupae (i.e., definitely last instar; 0.173 ± 0.006 mm, *n* = 35) bracket the lower and upper limit of the larval head width. Between these limits, an intermediate peak was detected around 0.135–0.140 mm in the histogram of larval head width (*n* = 590, bin width = 0.005 mm), suggesting the presence of an additional larval instar. Furthermore, the number and shape of larval hairs in these three head width ranges were clearly different; first instar: 100–150 simple smooth hairs, intermediate: 500–600 simple smooth hairs, last instar: 400–500 anchor-tipped hairs. According to the agreement between morphometrical and chaetotaxical features, we estimated that there are three instars of *M. triviale* worker larvae (Table 1). The larval head width of first (0.122 ± 0.006 mm, range: 0.107–0.137 mm, *n* = 84), second (0.137 ± 0.006 mm, range: 0.124–0.152 mm, *n* = 226) and third (0.175 ± 0.008 mm, range: 0.146–0.191 mm, *n* = 116) instar larvae overlapped (Fig. 1). The average linear growth increment between the larval instars was 1.20 (1.12 between the first and second instars; 1.27 between the second and third instars).

**FIGURE 1.**
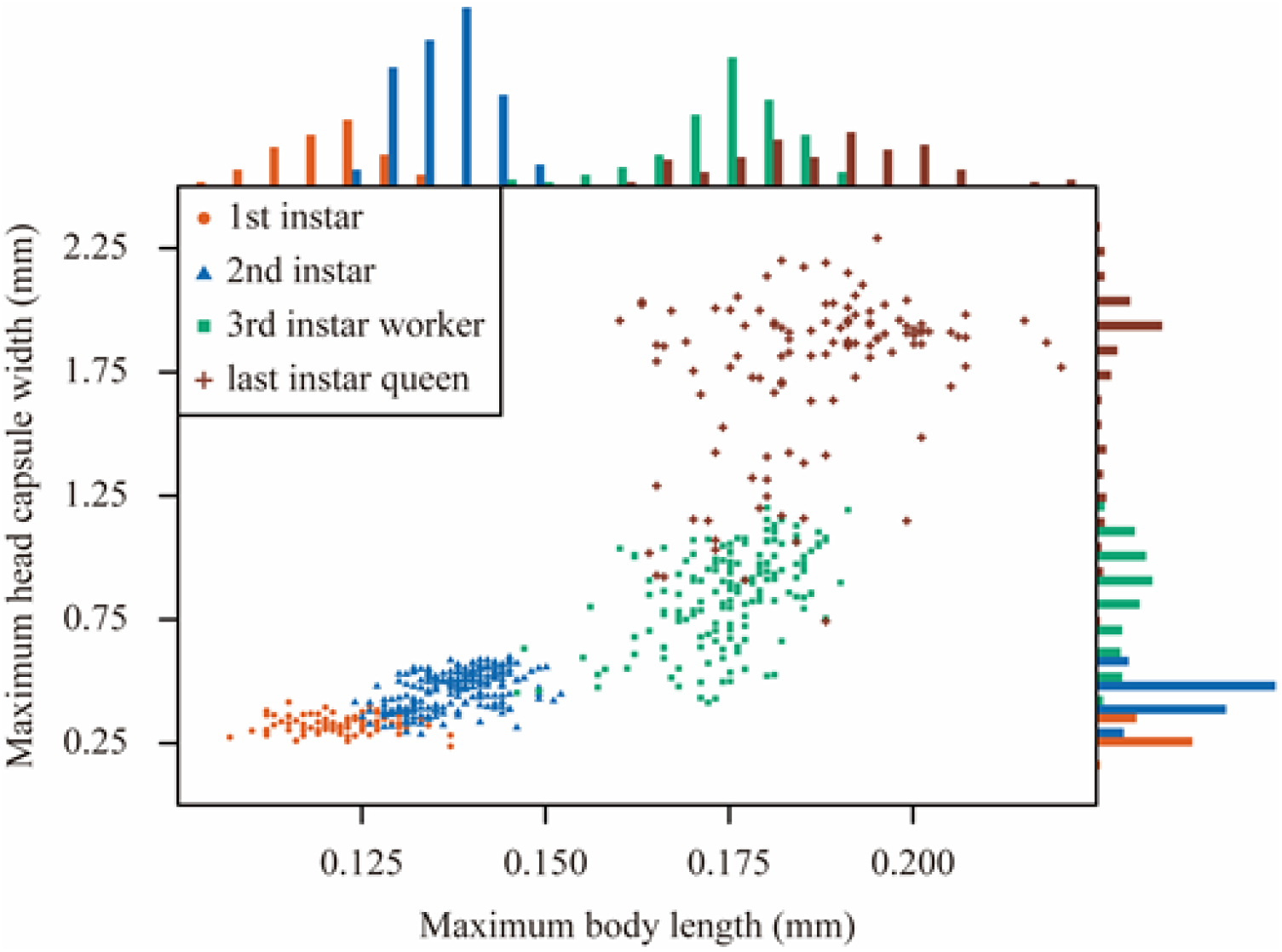
Relationship between the maximum head capsule width (mm) and the maximum body length (mm) of *Monomorium triviale* larvae. The 1st instar (orange circle, *n* = 84), 2nd instar (blue triangle, *n* = 226), 3rd instar worker (green square, *n* =164) and last instar queen (brown cross, *n* = 116) larvae were determined by morphology and chaetotaxy. Histogram bin width of head capsule width and body length is 0.005 mm and 0.1 mm respectively.

**Table 1.** Morphological and chaetotaxical characteristics in different castes and instars of *M. triviale*.

| **Larval instar** | **Head width**<br>**(mm)** | **Body length**<br>**(mm)** | **Hair**<br>**number** | **Hair type** | **Protuberances** |
| --- | --- | --- | --- | --- | --- |
| **1st instar** | 0.122 ± 0.006<br>(0.107–0.137) | 0.328 ± 0.036<br>(0.237–0.416) | 100–150 | simple<br>smooth | absent |
| **2nd instar** | 0.137 ± 0.006<br>(0.124–0.152) | 0.459 ± 0.074<br>(0.281–0.593) | 500–600 | simple<br>smooth | absent |
| **3rd instar**<br>**worker** | 0.175 ± 0.008<br>(0.146–0.191) | 0.847 ± 0.192<br>(0.414–1.20) | 400–500 | anchor-tipped | absent |
| **Last instar**<br>**queen** | 0.187 ± 0.013<br>(0.16–0.220) | 1.756 ± 0.329<br>(0.742–2.291) | almost<br>hairless | simple<br>smooth | present |

### Description of the immatures of *M. triviale*

#### Egg

The egg is oval in profile and presents a whitish translucent chorion (Fig. 2A). Length is 0. 31 ± 0. 02 mm, range 0.24–0.35 mm; width is 0.19 ± 0.01 mm, range 0.16–.022 mm (all *n* = 183). The length:width ratio for the species is 1.62 on average.

**FIGURE 2.**
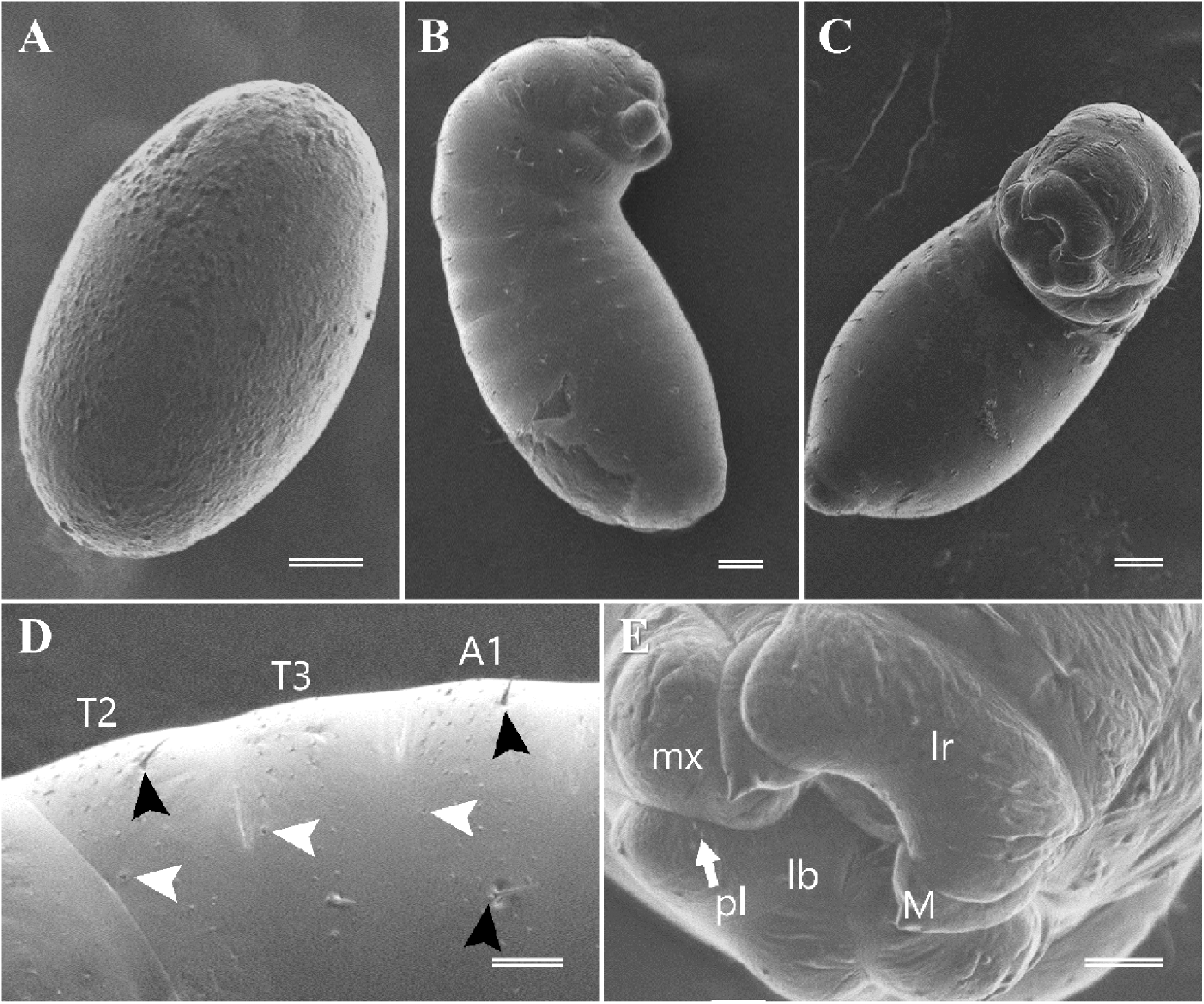
Egg and first instar larva of *Monomorium triviale*. **A.** side view of egg; **B.** habitus of first instar larva in lateral view; **C.**habitus of first instar larva in ventral view; **D.** lateral body surface of first instar larva, showing the spiracles (white arrowheads) and unbranched hairs (black arrowheads) on the thoracic (T2–T3) and abdominal (A1) segments; **E.** mouthparts of first instar larvae: labrum (lr), maxilla (mx), mandible (M), labium (lb) and labial palp (pl). Scale bars: 33.3 μm (A, B, C); 14.2 μm (D, E).

### First instar

#### Body

Whitish, profile ‘pheidoloid’ *sensu* Wheeler & Wheeler (1976) which is characterized by short and straight abdomen, short prothorax, and ends rounded to ventral direction (Fig. 2B); distinct anterior somites; head subcircular; anus subterminal (Fig. 2C). Body 0.328 ± 0.036 mm long (0.237–0.416 mm), 0.161 ± 0.014 mm wide (0.123–0.20 mm, *n* = 84). Length through spiracles 0.44 ± 0.34 mm (*n* = 6). Body hair varying in number (100–150; *n* = 5), unbranched, 8.4 ± 2.0 μm long (*n* = 25 of 5 individuals), distance between two adjacent hairs 25.6 ± 9.1 μm (*n* = 25 from 5 individuals, Fig. 2D). Body with 10 spiracles; distance between spiracles 29.2 ± 9.0 μm (*n* = 5). Spiracle opening unornamented; diameter of the first spiracle 0.8 ± 0.3, other spiracles 0.5 ± 0.2 μm (*n* = 5).

#### Head capsule

Cranium subcircular, 0.125 ± 0.011 mm in length (0.091–0.149 mm), 0.122 ± 0.006 mm in width (0.107–0.137 mm, *n* = 84); cranium index (length / width × 100; Bharti *et al*. 2019) = 102.3 ± 9.6. Head surface smooth with 26 unbranched hairs: 4 along the ventral border of the clypeus; 12 hairs on each gena; 4 on frons; 2 on vertex; and 4 along the occipital border. Length of hairs 8.7 ± 1.9 μm (*n* = 25 of 5 individuals).

#### Mouthparts

Clypeus clearly defined, with no sensilla (Fig. 2E); labrum bilobed, 62.1 ± 1.7 μm wide (*n* = 5), with six sensilla on the anterior surface. Mandibles subtriangular with broad base, blade ending in an apical tooth, lacking medial teeth (Fig. 2E), 0.034 ± 0.002 mm long (0.029–0.038 mm, *n* = 9). Maxilla conoidal in shape, 36.1 ± 2.1 μm wide (*n* = 10). Galea with two sensilla. Maxillary palpus a skewed peg with two sensilla. Labium elliptical, 52.3 ± 2.8 μm wide (*n* = 5). Labial palpus with a sensillum on the top.

### Second instar

#### Body

Whitish, profile pheidoloid in shape (Fig. 3A); distinct anterior somites; head subcircular; anus subterminal (Fig. 3B). Body about 0.459 ± 0.074 mm long (0.281–0.593 mm), 0.189 ± 0.021 mm wide (0.149–0.250 mm, *n* = 226). Length through spiracles 0.77 ± 0.18 mm (*n* = 5). Body hair varying in number (500–600; *n* = 5), unbranched, 6.8 ± 2.0 μm long (*n* = 25 of 5 individuals), distance between two adjacent hairs 13.8 ± 2.9 μm (*n* = 25 from 5 individuals, Fig. 3C). Body with 10 spiracles; distance between spiracles 46.8 ± 4.8 μm (*n* = 5). Spiracle opening unornamented; diameter of the first spiracle 2.1 ± 0.3, other spiracles 0.6 ± 0.1 μm (*n* = 5).

**FIGURE 3.**
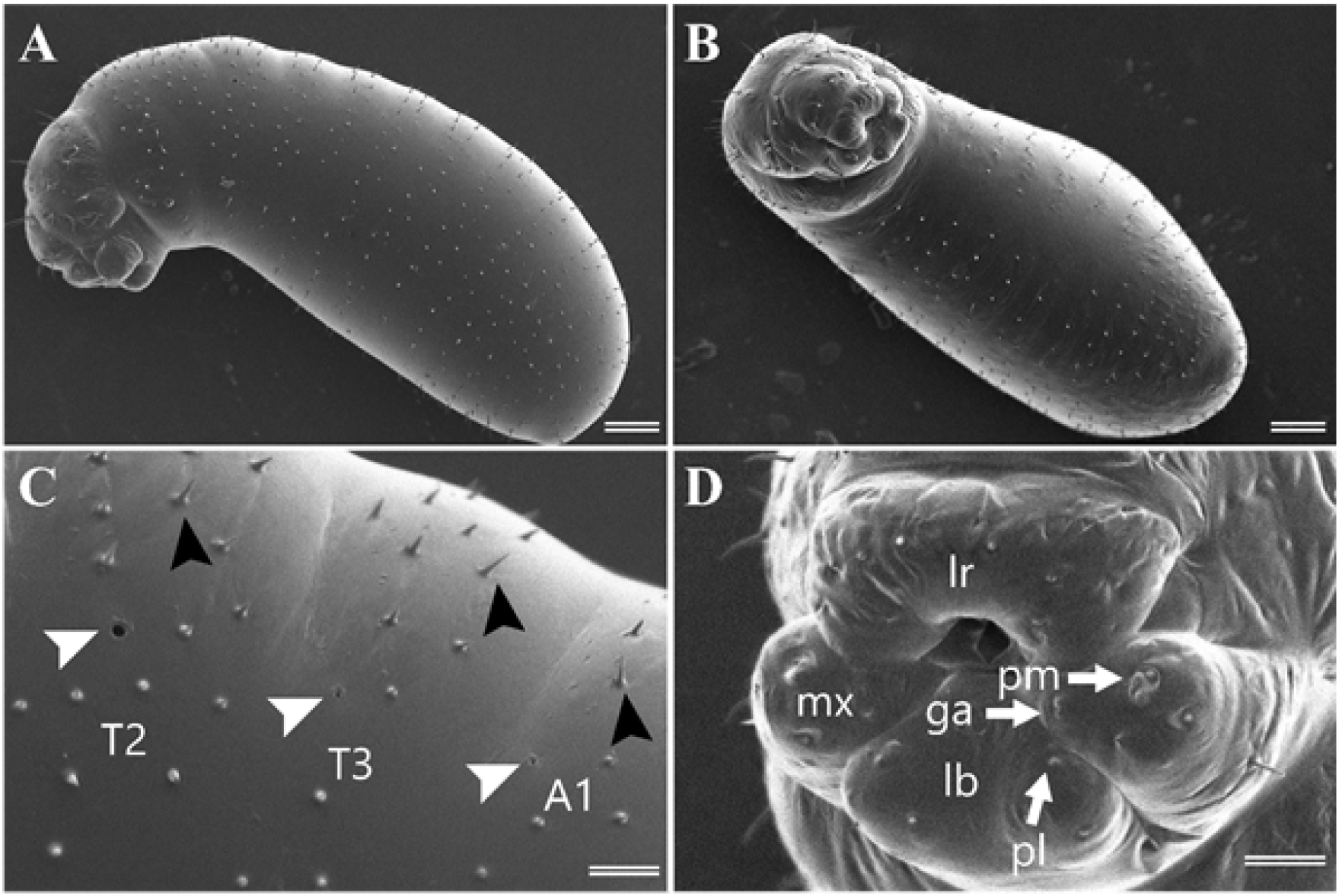
Second instar larva of *Monomorium triviale*. **A.** habitus in lateral view; **B.** habitus in ventral view; **C.** lateral body surface, showing the spiracles (white arrowheads) and unbranched hairs (black arrowheads) on the thoracic (T2–T3) and abdominal (A1) segments; **D.** mouthparts: labrum (lr), maxilla (mx), labium (lb), maxillary palp (pm) and labial palp (pl). Scale bars: 50 μm (A, B); 14.2 μm (C, D).

#### Head capsule

Cranium subcircular, 0.146 ± 0.016 mm in length (0.108–0.189 mm), 0.137 ± 0.006 mm in width (0.124–0.152 mm, *n* = 226); cranium index = 106.4 ± 10.5. Head surface smooth with 26 unbranched hairs: 4 along the ventral border of the clypeus; 12 hairs on each gena; 4 on frons; 2 on vertex; and 4 along the occipital border. Length of hairs 12.9 ± 3.3 μm (*n* = 25 of 5 individuals).

#### Mouthparts

Clypeus clearly defined, with no sensilla; Labrum bilobed, 67.3 ± 1.3 μm wide (*n* = 5, Fig. 3D), with six sensilla on the anterior surface. Mandibles ‘ectatommoid’ as defined by Wheeler & Wheeler (1976): “Subtriangular; with a medial blade arising from the anterior surface and bearing one or two medial teeth; apex curved medially to form a tooth”, sclerotized and bearing at least two teeth, 0.042 ± 0.002 mm long (0.039–0.046 mm, *n* = 13). Maxilla conoidal in shape, 40.1 ± 1.6 μm wide (*n* = 8). Galea with two sensilla. Maxillary palpus a skewed peg with two sensilla. Labium elliptical, 51.5 ±2.5 μm wide (*n* = 5). Labial palpus with a sensillum on the top.

### Third instar

#### Body

Whitish, profile pheidoloid in shape (Fig. 4A); distinct anterior somites (Fig. 4B); head subcircular (Fig. 4C); anus subterminal. Body about 0.847 ± 0.192 mm long (0.414–1.20 mm), 0.360 ± 0.087 mm wide (0.192–0.608 mm, *n* = 164). Length through spiracles 0.95 ± 0.05 mm (*n* = 5). Body hair varying in number (400–500; *n* = 5), deeply branched, 32.5 ± 4.4 μm long (*n* = 25 of 5 individuals), distance between two adjacent hairs 26.4 ± 5.3 μm (*n* = 25 from 5 individuals, Fig. 4D). Body with 10 spiracles; distance between spiracles 57.8 ± 9.2 μm (*n* = 5). Spiracle opening unornamented; diameter of the first spiracle 2.2 ± 0.4, other spiracles 0.7 ± 0.2 μm (*n* = 5, Fig. 4E).

**FIGURE 4.**
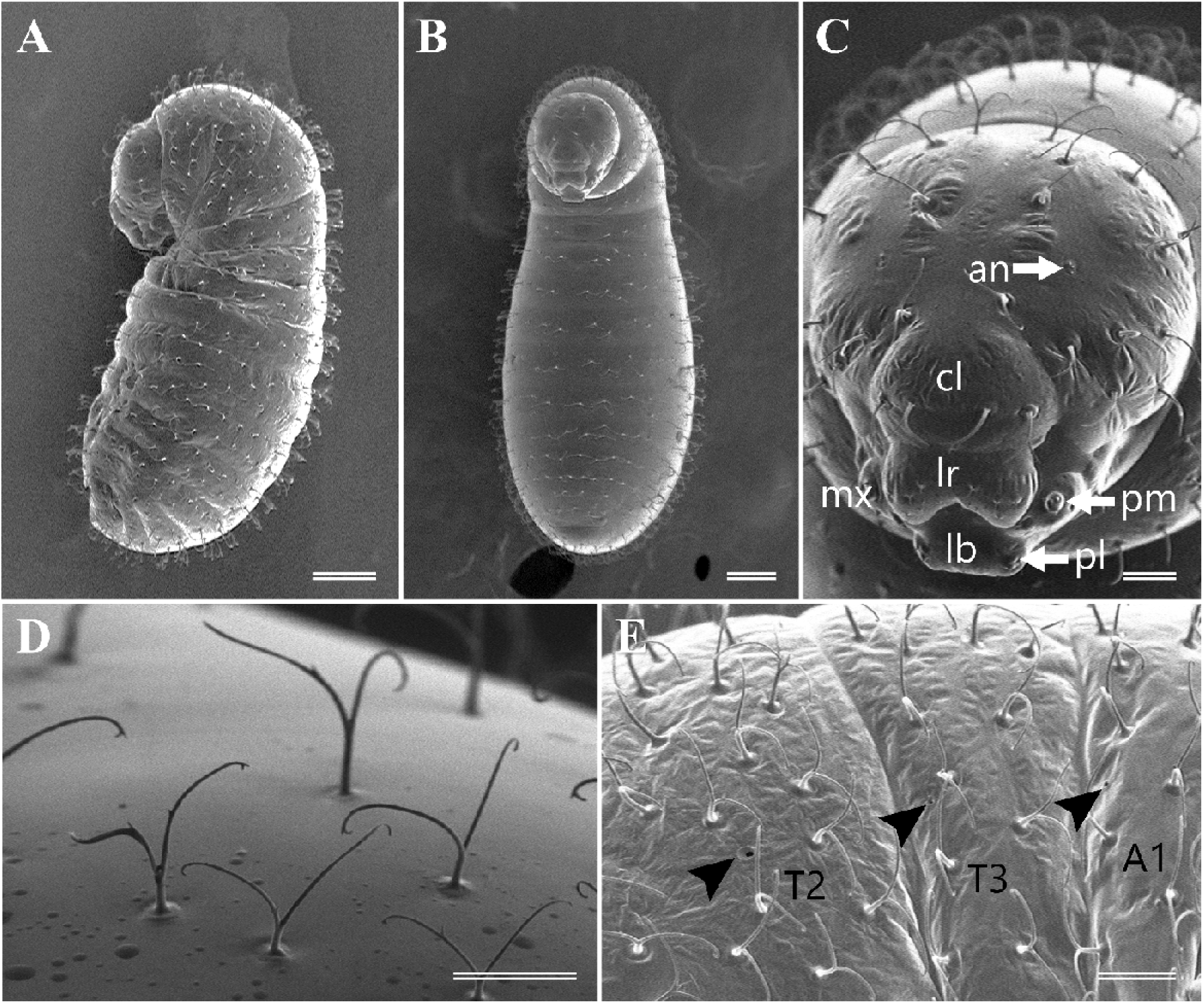
Third instar larva of *Monomorium triviale*. **A.** habitus in lateral view; **B.** habitus in ventral view; **C.** head and mouthparts: antenna (an), clypeus (cl), labrum (lr), maxilla (mx), labium (lb), maxillary palp (pm) and labial palp (pl); **D.** anchor-shaped body hairs on dorsal thoracic somite; **E.** lateral body surface, showing the spiracles (black arrowheads) on the thoracic (T2–T3) and abdominal (A1) segments. Scale bars: 100 μm (A, B); 25 μm (C, D, E).

#### Head capsule

Cranium subcircular, 0.189 ± 0.017 mm in length (0.134–0.227 mm), 0.175 ± 0.008 mm in width (0.146–0.191 mm, *n* = 164); cranium index = 108.3 ± 8.0. Head surface smooth with Twenty-six branched hairs: 4 along the ventral border of the clypeus; 12 hairs on each gena; 4 on frons; 2 on vertex; and 4 along the occipital border. Length of hairs 29.1 ± 4.7 μm (*n* = 25 of 5 individuals).

#### Mouthparts

Clypeus clearly defined, with no sensilla (Fig. 4D); Labrum bilobed, 65.5 ± 1.3 μm wide (*n* = 5), with six sensilla on the anterior surface. Mandibles ectatommoid, with three distinct teeth, more sclerotized than those of the second instar, 0.067 ± 0.003 mm long (0.063–0.071 mm, *n* = 5). Maxilla conoidal in shape, 46.9 ± 1.2 μm wide (*n* = 8). Galea with two sensilla. Maxillary palpus a skewed peg with two sensilla. Labium elliptical, 55.5 ± 4.2 μm wide (*n* = 5). Labial palpus with a sensillum on the top.

### Last instar queen

#### Body

Whitish, profile ‘aphaenogastroid’ *sensu* Wheeler & Wheeler (1976) which is characterized by constriction between thorax and abdomen (Fig. 5A); distinct anterior somites; head subcircular; anus subterminal (Fig. 5B). Body about 1.756 ± 0.329 mm long (0.742–2.291 mm), 0.873 ± 0.194 mm wide (0.408–1.214 mm, *n* = 116). Length through spiracles 1.8–2.1 mm (*n* = 2). Abdominal somites hairless (Fig 5A-C), prothoracic body hair varying in number 50–70; *n* = 2), unbranched, 52.8 ± 14.9 μm long (*n* = 20 of 2 individuals, Fig. 5D, E), distance between two adjacent hairs 25.3 ± 5.9 μm (*n* = 20 from 2 individuals). Body with 10 spiracles; distance between spiracles 118.5–213.4 μm (*n* = 1). Spiracle opening unornamented; diameter of the first spiracle 2.7–3.1, other spiracles 1.3–1.8 μm (*n* = 2).

**FIGURE 5.**
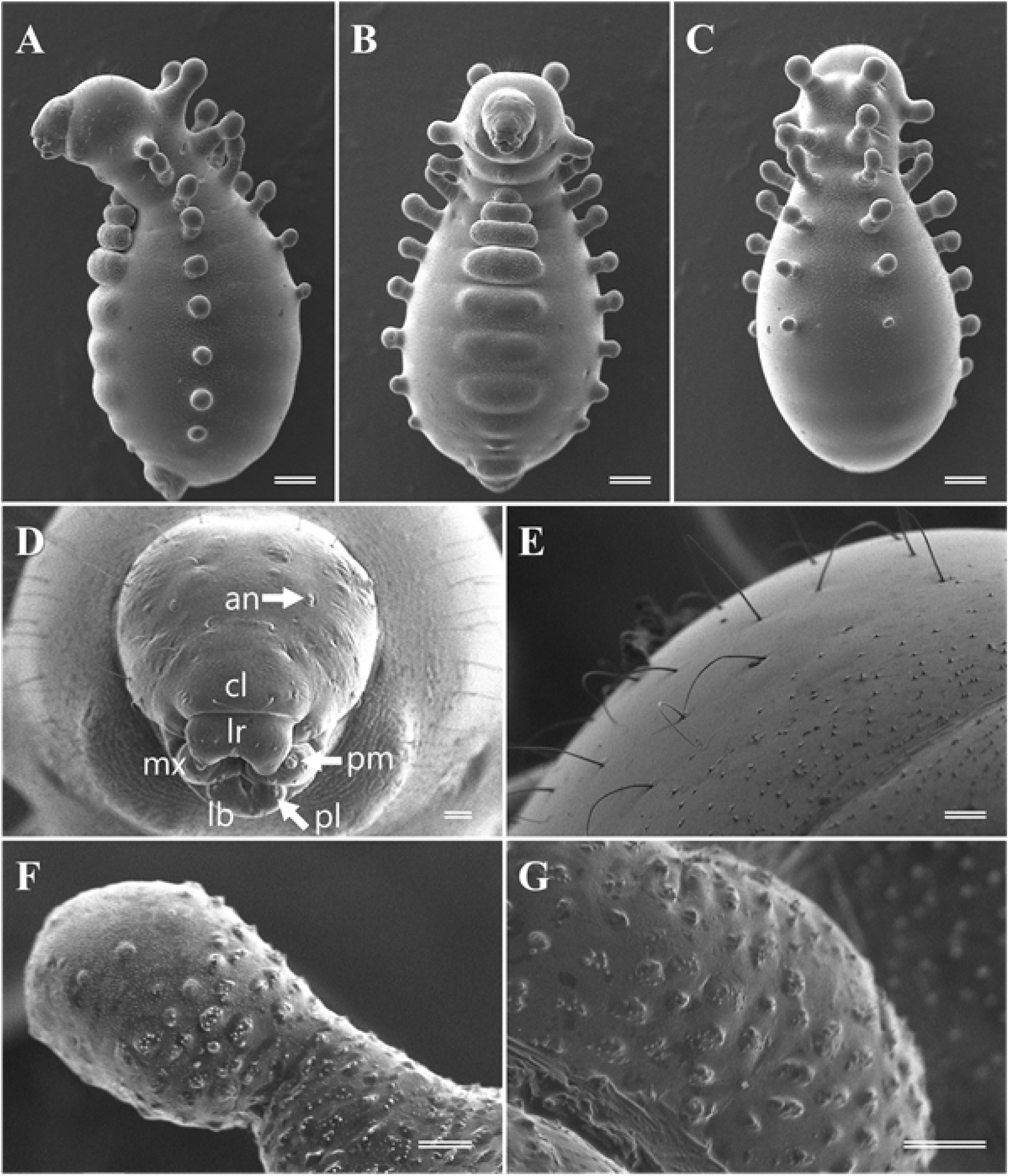
Last instar larva of *Monomorium triviale* queen. **A.** habitus in lateral view; **B.** habitus in ventral view; **C.** habitus in dorsal view; **D.** head and mouthparts: antenna (an), clypeus (cl), labrum (lr), maxilla (mx), labium (lb), maxillary palp (pm) and labial palp (pl); **E.** body hairs; **F.** doorknob-like tubercle; **G.** mid-ventral boss. Scale bars: 142 μm (A, B, C); 20 μm (D, E, F, G).

#### Head capsule

Cranium subcircular (Fig. 5D), 0.194 ± 0.027 mm in length (0.096–0.256 mm), 0.187 ± 0.013 mm in width (0.16–0.220 mm, *n* = 116); cranium index = 103.9 ± 12.7. Head surface smooth with Twenty-six unbranched hairs: 4 along the ventral border of the clypeus; 12 hairs on each gena; 4 on frons; 2 on vertex; and 4 along the occipital border. Length of hairs 28.3 ± 9.3 μm (*n* = 20 of 2 individuals).

#### Mouthparts

Clypeus clearly defined, with no sensilla; Labrum bilobed, 77.5 ± 2.2 μm wide (*n* = 5), with six sensilla on the anterior surface. Mandibles ectatommoid, with 3 teeth, 0.069–0.070 mm long (*n* = 2). Maxilla conoidal in shape, 51.8 ± 3.4 μm in wide (*n* = 8). Galea with two sensilla. Maxillary palpus a skewed peg with two sensilla. Labium elliptical, 59.1 ± 2.2 μm wide (*n* = 5). Labial palpus with a sensillum on the top.

#### Protuberances

Total 37 protuberances; 9 pairs on dorsal, 6 pairs on lateral and unpaired 7 on mid-ventral (Table 2). Paired doorknob-like dorsal tubercles on 2–3rd thoracic and 1–4th abdominal somites, 72.3 ± 9.4 μm diameter, 134.0 ± 52.7 μm long (*n* = 12). Lateral doorknob-like tubercles on 2–3rd thoracic and 1–7th abdominal somites, 73.4 ± 9.9 μm diameter, 103.7 ± 33.9 μm long (*n* = 36). Ventral protuberances “bosses” as defined by Wheeler & Wheeler (1976); “an elevated structure with a rounded terminus” in shape. Anterior four bosses on the 3rd thoracic and 1–3rd abdominal somites clearly defined by outlines, 260.7 ± 95.0 μm wide, 105.4 ± 40.5 μm long, 116.5± 35.9 μm high (*n* = 8). Posterior three bosses on the 4–6th abdominal somites slightly raised from body without distinct border. All protuberances surface uneven with minute papillae (Fig. 5F, G). We did not find any opening or duct-like structures on the surface of protuberances.

**TABLE 2.** Protuberance position and size in the last instar queen larvae of *Monomorium triviale*.

|  | Lateral tubercle size<br>(µm) |  | Dorsal tubercle size<br>(µm) |  | Ventral boss size (µm) |  |  |
| --- | --- | --- | --- | --- | --- | --- | --- |
| Somites | diameter | length | diameter | length | width | length | height |
| Thoracic 1 |  | - |  | - |  | - |  |
| Thoracic 2 | 61-76 | 116-141 | 80-95 | 184-210 |  | - |  |
| Thoracic 3 | 77-87 | 102-120 | 67-82 | 160-210 | 101-217 | 52-75 | 43-79 |
| Abdominal 1 | 60-86 | 130-161 | 70-73 | 135-173 | 181-277 | 66-109 | 116-122 |
| Abdominal 2 | 66-93 | 119-168 | 69-73 | 113-135 | 238-349 | 89-138 | 123-152 |
| Abdominal 3 | 68-92 | 107-127 | 62-69 | 82-95 | 293-430 | 134-180 | 137-159 |
| Abdominal 4 | 73-82 | 92-111 | 59-67 | 46-65 |  | indistinct |  |
| Abdominal 5 | 66-81 | 79-90 |  | - |  | indistinct |  |
| Abdominal 6 | 55-75 | 54-70 |  | - |  | indistinct |  |
| Abdominal 7 | 55-66 | 38-58 |  | - |  | - |  |
| Abdominal 8 |  | - |  | - |  | - |  |
| Abdominal 9 |  | - |  | - |  | - |  |
| Abdominal 10 |  | - |  | - |  | - |  |

### Pupae

Exarate, with no cocoon, 0.99–1.12 mm long in workers (*n* = 10, Fig. 6A) and 2.20–2.49 mm in queens (*n* = 2, Fig. 6B), milky-white when young; eyes becoming black and body gradually darkened to a yellowish-brown as they develop into imagos. Queen wings vestigial (i.e., brachypterous, Fig. 6C).

**FIGURE 6.**
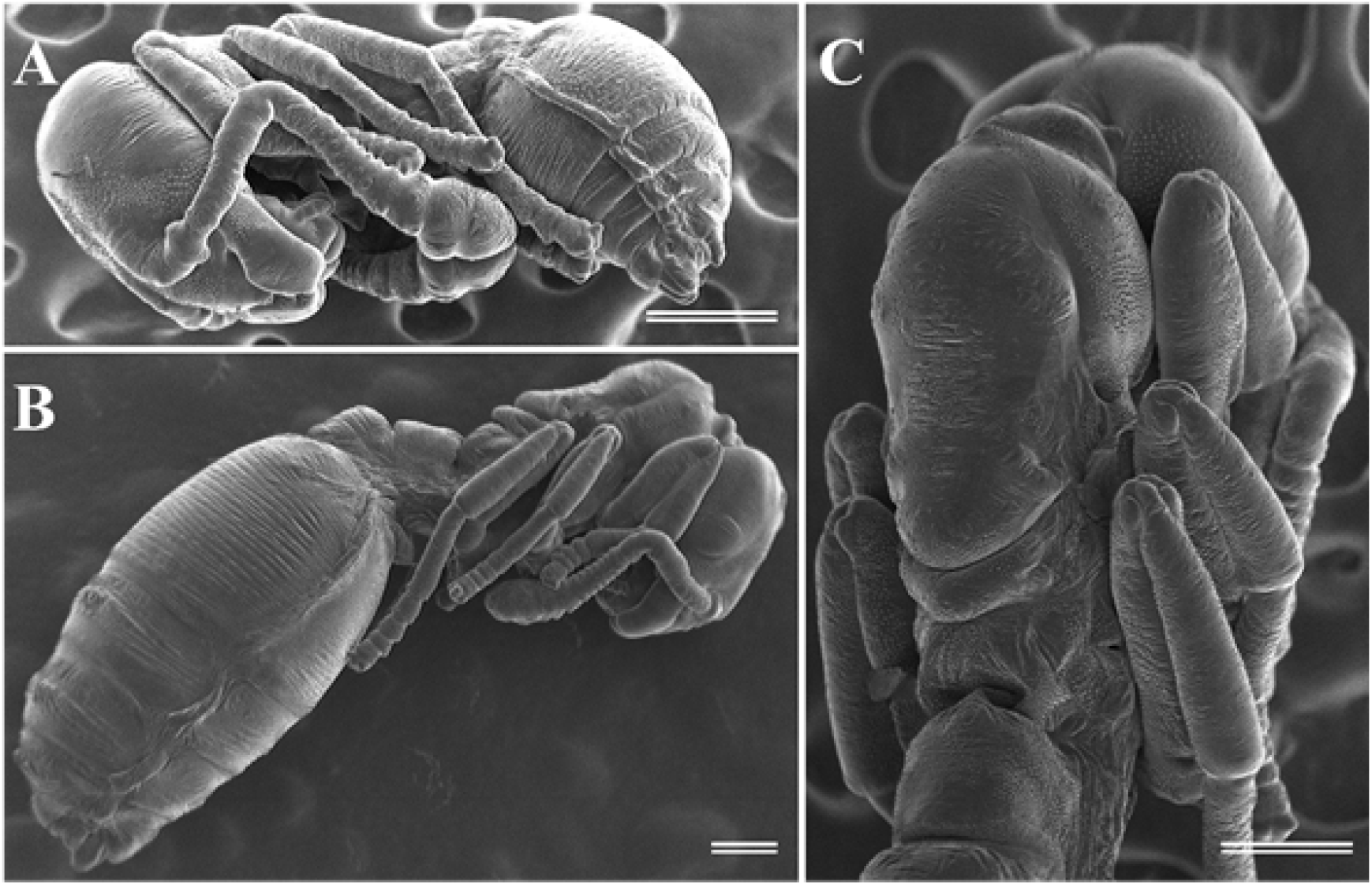
Pupae of *Monomorium triviale*. **A.** habitus of worker pupa in lateral view; **B.** habitus of queen pupa in lateral view; **C.** thorax of queen pupa in dorsal view. All scale bars = 200 μm.

### Caste dimorphism in congeneric species

The last instar larvae of the three observed *Monomorium* species, *M. hiten, M. intrudens* and *M. chinense* shared distinct anterior somites, subcircular head shape and subterminal anus. The worker larvae of all three species indicated the pheidoloid shape in profile and were covered with anchor-shaped hairs on the thoracic and abdominal somites (Fig. 7A–C). In *M. hiten*, the larvae of reproductive males and females (= queens) also had pheidoloid shape (Fig. 7D). However, in *M. intrudens* (Fig. 7E) and *M. chinense* (Fig. 7F), the reproductive larvae indicated “attoid” shape as defined by Wheeler & Wheeler (1976); “diameter approximately equal to distance from labium to anus” in profile. The anchor-shaped hairs were not found in the reproductive larvae of all three species.

**FIGURE 7.**
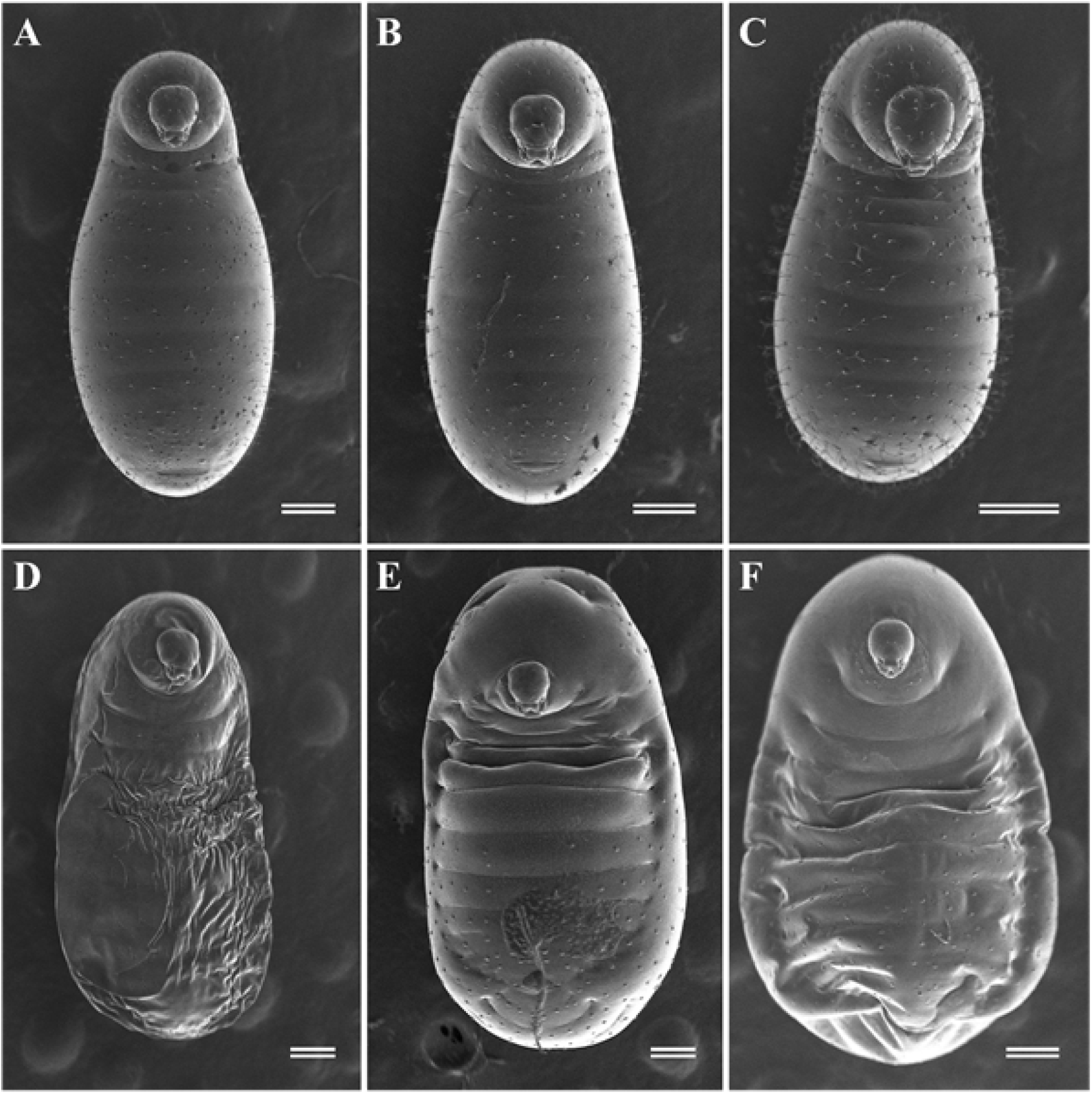
Habitus of last instar *Monomorium* larvae in ventral view. **A.** *M. hiten* worker; **B.** *M. intrudens* worker; **C.** *M. chinense* worker; **D.** *M. hiten* reproductive; **E.** *M. intrudens* reproductive; **F.** *M. chinense* reproductive. All scale bars = 200 μm.

### Internal morphology of last instar queen larvae

Histological observations of the queen larva showed that the dorsal and lateral doorknob-like tubercles (Fig. 8A, B) and ventral bosses (Fig. 8C, D) were formed by extended epidermis and cuticle. There are no muscles or innervation inside the dorsal and lateral tubercles and ventral bosses. The thickness of the cuticle and epidermis was 8.6 ± 1.8 μm (*n* = 7) and 11.6 ± 3.3 μm (*n* = 7) in the doorknob-like tubercles, two times thicker than the other part of the larval body, which was 4.2 ± 0.2 μm (*n* = 9) and 4.9 ± 1.0 μm (*n* = 9). The ventral bosses were formed by slightly thicker cuticle 5.78 ± 1.11 μm (*n* = 5) and markedly thicker epidermis 16.42 ± 4.09 μm (*n* = 5). In the ventral bosses, the small cells (possibly fat bodies) surrounded by epidermis were observed (Fig. 8D). We did not find any specialized cells and duct-like structures in the doorknob-like tubercles and in the ventral bosses.

**FIGURE 8.**
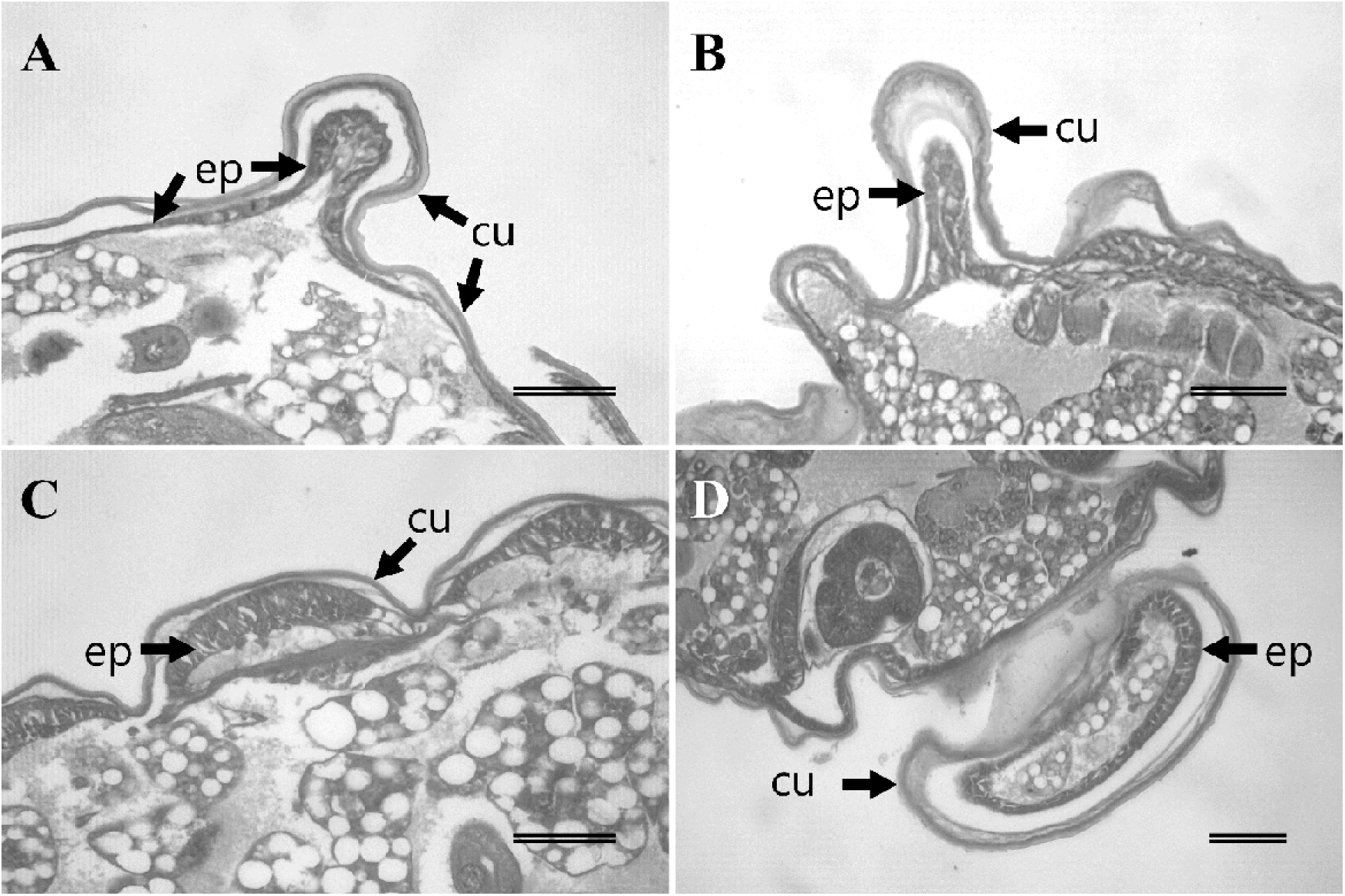
Histological section through the protuberances of last instar queen larvae, showing epidermis (ep) and cuticle (cu). **A.** cross section of dorsal doorknob-like tubercles; **B.** cross section of lateral doorknob-like tubercles; **C.** longitudinal section of mid-ventral bosses; **D.** cross section of mid-ventral bosses. All scale bars = 50 μm.

### Developmental periods

During the 53-day experimental period, 211 eggs and 166 first instar larvae appeared in the artificial nests (Table 3). The average duration between oviposition and hatching was 13.39 ± 2.28 days. Out of 70 induced eggs, 32 young (= first and second instar) larvae hatched and all successfully turned into the last instar larvae in an average of 7.31 ± 1.00 days. Seventy-two of 80 induced young larvae molted to the last instar larvae and 51 of them turned into prepupae in 18.31 ± 3.29 days, and all molted to pupae in 4.43 ± 1.10 days. Thirty-three of 70 induced last instar larvae turned to pupae and 22 individuals successfully emerged as worker imago in 16.87 ± 1.55 days. In total, the overall developmental period of *M. triviale* worker was estimated to be 59 days.

**TABLE 3.** The developmental periods of the immature stages of *M. triviale* and the number of observed individuals for each stage.

| Developmental stage | Source nests | The numbers of individuals |  |  | Developmental period (day) |  |
| --- | --- | --- | --- | --- | --- | --- |
| | | Total observed | Turned to target stage | Molted to next stage | mean ± SD | range |
| Egg | 18 | NA | 211 | 166 | 13.39 ± 2.28 | 8-22 |
| Young larva | 7 | 70 | 32 | 32 | 7.31 ± 1.00 | 6-9 |
| Old larva | 8 | 80 | 72 | 51 | 18.31 ± 3.29 | 13-26 |
| Prepupa | 8 | 51 | 51 | 51 | 4.43 ± 1.10 | 1-7 |
| Pupa | 7 | 70 | 33 | 23 | 16.87 ± 1.55 | 13-20 |

## Discussion

Based on the combination of the body size and body hair features, three larval instars were estimated for *M. triviale*. This is in general the lowest number of larval instars in Formicidae and has been reported from four subfamilies: Dorylinae, Ectatomminae, Formicinae and Myrmicinae (Solis *et al*. 2010a). In Myrmicinae, three larval instars were known from 16 species including the congeners, *M. pharaonis* and *M. floricola* (Solis *et al*. 2010a; Bharti & Gill 2011; Bharti *et al*. 2019). In *M. triviale*, calculated growth rates between the instars are consistent with the Dyar principle of a constant ratio of growth by molting between 1.1 and 1.9 (Parra & Haddad, 1989). The growth rate during the larval period of *M. triviale* was 1.19, which is smaller than that of *M. floricola* (1.23; Solis *et al*. 2010a) and *M. pharaonis* (1.36; Alvares *et al*. 1993). This difference may reflect the difference in body size of adult workers, ca. 1.5 mm for *M. triviale*, 1.7–2.0 mm for *M. floricola* (Wetterer 2010a), and 2.2–2.4 mm for *M. pharaonis* (Wetterer 2010b).

The total developmental period of *M. triviale* was estimated to be about 59 days (egg: 13 days + larva: 29 days + pupa: 17 days). These results are slightly longer than those of the tropical congeners *M. pharaonis* (egg: 11 days + larva: 22 days + pupa: 12 days, 27°C; Pontieri *et al*. bioRxiv) and *M. hiten* (egg: 12 days + larva: 17 days + pupa: 20 days, 24°C; Ito *et al*. 2021). It should be noted that the larval period of *M. triviale* may be extended under natural conditions because the larvae can stop their development and overwinter in the temperate zone of Japan.

Adams *et al*. (2021) suggested that the combination of quantitative and binary traits is effective in distinguishing the sex and caste of ant larvae. Our results showed that this approach is also viable in *M. triviale*. Although the body size of each larval instar overlapped (Fig.1), three larval instars could be separated by the number and shape of their body hairs (Table 1). The larvae of some *Monomorium* species have deeply branched anchor-shaped hairs (Solis *et al*. 2010a; Penick *et al*. 2012). In *M. triviale*, the third instar worker larvae possess these anchor-shaped hairs all over their body. On the other hand, the last instar larvae of the queen have only a few simple hairs on the prothorax. Moreover, the three congeneric species *M. intrudens, M. hiten* and *M. chinense* also showed the same caste-specific pattern: worker larvae had branched hairs, but reproductive larvae were almost hairless. Such caste dimorphism of body hairs was previously known in *M. minimum* (Wheeler & Wheeler 1955) and *M. pharaonis* (Edwards 1991). The absence of anchor-tipped hairs on the last instar could be a useful trait to distinguish the reproductive larvae of *Monomorium* species.

The caste dimorphism in the last larval instar of *M. triviale* is more striking than in other observed congeneric species. The queen larvae of *M. triviale* have “aphaenogastroid” shape in profile, the lateral and dorsal rows of doorknob-like tubercles on thoracic and abdominal segments and semi-elliptical protrusion on ventral abdominal segments. The distinct segmental patterns of these structures among lateral, dorsal and ventral sides, especially their absence on the first thoracic segment and the terminal abdominal segments (Table 2), might provide a good opportunity to investigate the caste-specific developmental processes.

Unusual forms of ant larvae have been described from diverse subfamilies (Table 4). In the Myrmicinae, the caste dimorphism similar to that in *M. triviale* has been reported in *Crematogaster* species. Some late-instar larvae of *C. rivai* var. *luctuosa* and *C. scutellaris* have rows of lateral protuberances on the abdominal segments (Menozzi 1930; Eidmann 1926), and it is speculated that these larvae differentiate into queens (Casevitz-Weulersse 1984). Contrastingly, in some genera of Dolichoderinae, protuberances have been found only on worker larvae (e. g. Solis *et al*. 2010b). Although their developmental homology remains unknown, this phylogenetic distribution would be suggestive of their evolutionary convergence.

**TABLE 4.**
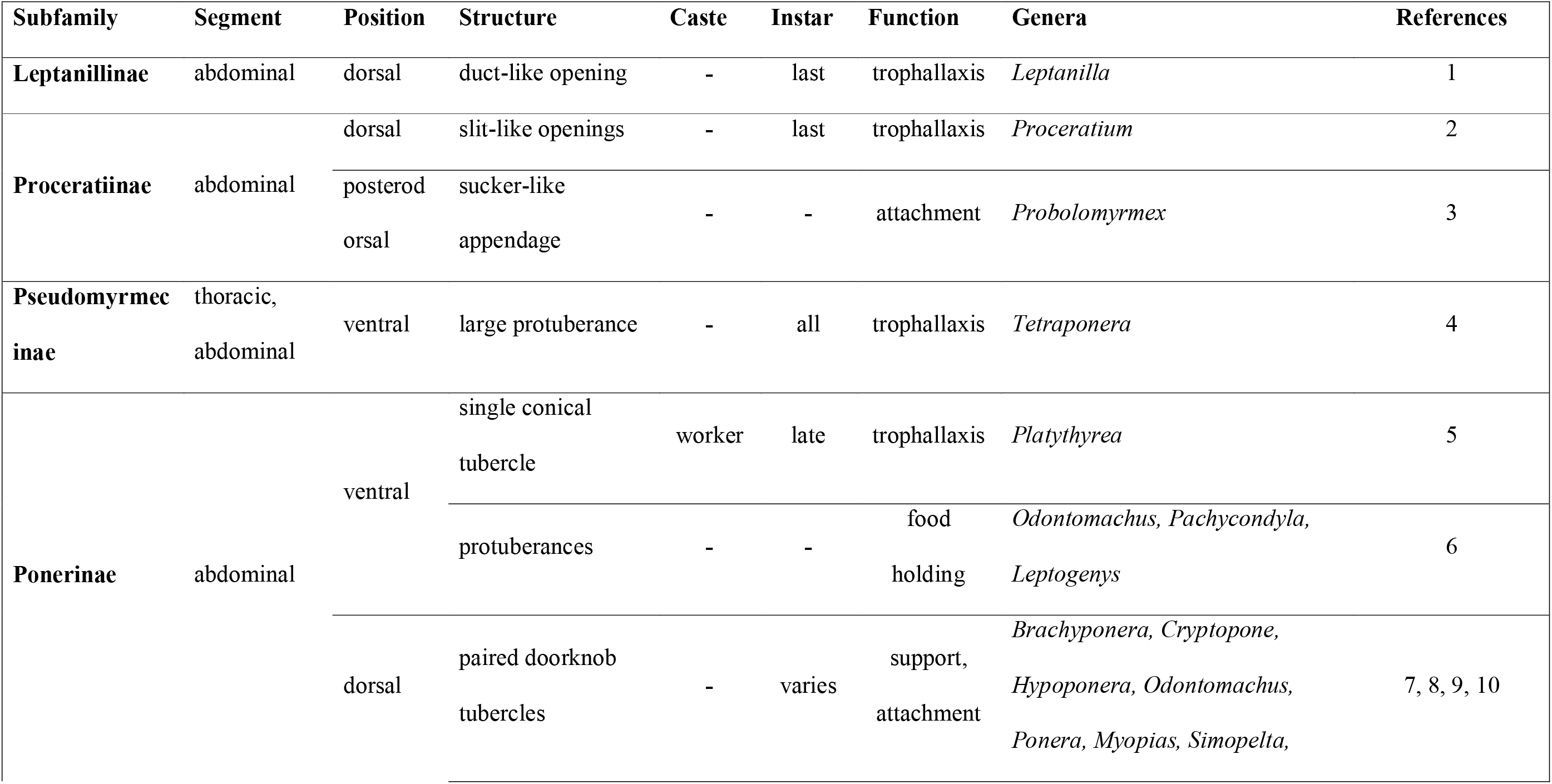
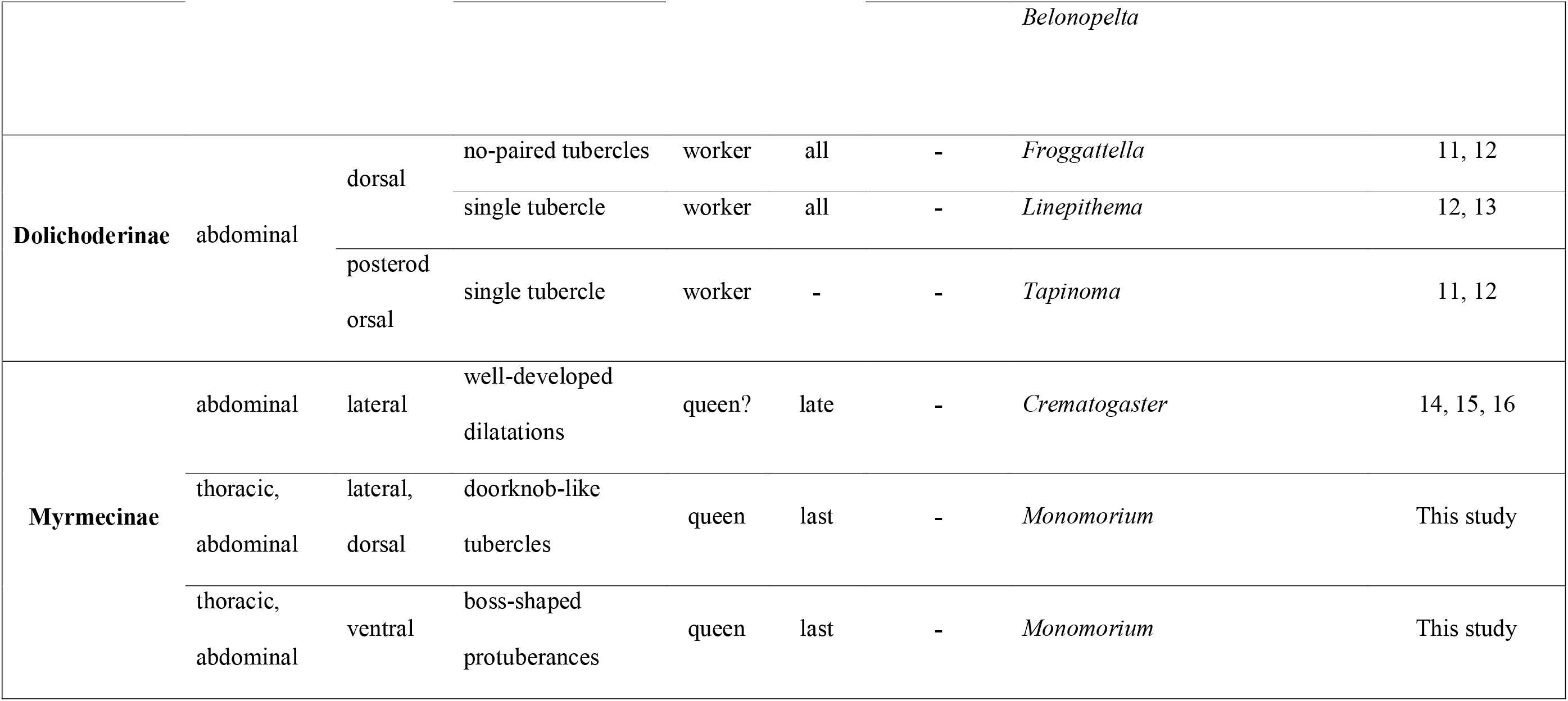
Previously reported unusual larval structures. References cited: 1, Masuko 1989; 2, Masuko 2019; 3, Taylor 1965; 4, Wheeler 1918; 5, Villet *et al*. 1990; 6, Petralia & Vinson; 1979; 7, Taylor 1967; 8, Wheeler & Wheeler 1971; 9, Peeters & Hölldobler 1992; 10, Fox *et al*. 2017; 11, Wheeler & Wheeler 1966; 12, Shattuck 1992; 13, Solis *et al*. 2010b; 14, Eidmann 1926; 15, Menozzi 1930; 16, Casevitz-Weulersse 1984.

Wheeler & Wheeler (1976) proposed five possible functions of protuberances in the ant larvae: (1) support of the body position and direction, (2) defense against cannibalism among larvae, (3) attachments to ceilings and walls of the nest, (4) organs for trophallaxis with adult workers, and (5) structures for holding food on the body surface. These hypotheses have not been sufficiently tested, with only a few exceptions such as Masuko (2019) and Fox *et al*. (2017). Our morphological and histological observations could not find any specialized structures such as duct-like openings and secretory cells in the protuberances of *M. triviale* queen larvae. As in many other cases, the function of queen-specific tubercles of the *M. triviale* larvae is still unclear at this time. Behavioral observations of the interaction between the workers and the queen larvae are required in the future. Further examination of the function of these unusual structures of larvae will help us understand the hidden but essential roles larvae play in complex ant societies.

## Acknowledgments

We are grateful to Kenji Matsuura who allowed us to use his laboratory. We thank Fuminori Ito for providing his *M. hiten* nest, Tomoya Kamada for his help during histological observation, and Matthew Kamiyama for his English corrections of this manuscript. Finally, we also thank Christian Peeters and Keiichi Masuko for their valuable comments on larval protuberances. This work was supported by a Japan Society for the Promotion of Science (JSPS) Research Fellowship for Young Scientists to NI (19J22242), JSPS KAKENHI Grant Number JP20K06080 to AG and a grant from the Secom Science and Technology Foundation to SD.

